# Effects of DNA extraction, DNA integrity, and laboratory on the precision of qPCR-based telomere length measurement - a multi-lab impartial study

**DOI:** 10.1101/2022.12.14.520438

**Authors:** Jue Lin, Simon Verhulst, Camilo Fernandez Alonso, Casey Dagnall, Shahinaz Gadalla, Waylon J. Hastings, Tsung-Po Lai, Idan Shalev, Ying Wang, Yun-Ling Zheng, Elissa Epel, Stacy Drury, the Telomere Research Network

## Abstract

Measuring telomere length (TL) with high precision is challenging. Currently there is insufficient understanding of the causes of variation in measurement precision, particularly for qPCR-based measurement. To better understand how DNA extraction protocols and laboratory-specific analytical factors influence qPCR-based TL measurement precision, we conducted a multi-laboratory study involving four labs and six DNA extraction protocols assaying the same blinded human whole blood samples. DNA extraction protocols differed in underlying principles (magnetic beads, salting out, silica membrane) and commercial kits. A fifth lab performed Telomere Restriction Fragment (TRF) analysis using Southern Blot technique with one DNA extraction protocol. All labs performed TL measurement using their standard procedures on two sets of fifty double blinded samples. Data was sent to a central point for unblinding and statistical analyses. Precision was quantified using the Intraclass Correlation Coefficient (ICC). Correlations with TRF measurements were also calculated. Repeated qPCR-based measurements of the same DNA extraction yielded ICC values ranging from 0.24 to 0.94. ICC values calculated over measurements of repeated DNA extractions were on average 0.23 lower and ranged from 0.02 to 0.83. The latter ICC estimates more strongly predicted the association between qPCR- and Southern blot-based measurements across the protocol / lab combinations (R^2^=0.56 vs. R^2^=0.93). We conclude that ICC calculated over measurements on repeated DNA extractions reliably indicates measurement precision, while ICC calculated over multiple measurements of the same DNA extraction overestimates measurement precision. Variation in ICC was driven by variation between laboratories, with few consistent DNA extraction protocol effects. Values of DNA integrity and purity generally characterized as reflecting high sample quality, (e.g. OD 260/280 of 1.8 and OD 260/230 of 2.0) were associated with qPCR-based measurement precision, but did not always predict higher ICCs.

## Introduction

The number of studies reporting on telomere length (TL) as a biomarker of life course exposures and a predictor of health outcomes has exponentially increased over the last two decades (1). Telomeres have generated broad interest due to their high conservation across vertebrate species and their association with processes across life history and outcomes (2, 3). Composed of tandem DNA repeated sequences, proteins, and RNA, telomeres fundamentally ensure chromosomal integrity, prevent DNA damage, and, when telomeres shorten to a critical length, may trigger cellular apoptosis, senescence, or terminal differentiation. Further, an increasing literature-base has associated telomere dynamics with regulation of gene expression across the chromosome and DNA damage responses highlighting the complex influence of telomere dynamics on biology, cellular function, and aging (4–7). Despite this enthusiasm and a transdisciplinary interest, accuracy and precision in TL measurements have been a persistent concern. Inconsistency in TL associations with health outcomes, environmental exposures, and psychosocial stress, potentially due to variability of protocols and methodologic reporting, indicated a need for a critical evaluation of TL assay precision, and the factors that impact measurement precision, to ensure the scientific rigor in this multi-disciplinary field (8–11). To assist in this evaluative process, the National Institute of Environmental Health Sciences and National Institute on Aging invested in the creation of a Telomere Research Network (TRN) (https://trn.tulane.edu/). The TRN is focused on addressing key methodologic questions in telomere length measurement and facilitation of broader discussions related to the utilization of TL as an indicator of psychosocial and environmental exposure and a predictor of disease across the life course.

A key directive of the TRN was to implement scientifically rigorous cross-method comparisons and refine the understanding of the factors influencing TL measurement precision within laboratories, across laboratories, across different methods (e.g. qPCR, fluorescence in situ, Southern blot, mixed methods) and across biological sample types applicable to population based studies. We utilize the term precision as equivalent to reproducibility defined by the National Academy of Science as “the obtaining of consistent results using the same data, methods, and conditions of analysis.” (12). As a first step in increasing the precision of TL studies, and consistent with the recommendation 4.1 of the National Academy of Science Reproducibility and Replicability in Science guidelines, the TRN developed minimum reporting recommendations for qPCR-based TL measurement (<u>doi: 10.31219/osf.io/9pzst</u>). This collaborative group further determined a critical need to evaluate, in a systematic manner, the impact of DNA extraction on assay precision within and across laboratories. Several previous studies have examined variability in TL measurement precision (2, 13–18). Universally, these studies have focused on the impact of DNA extraction methodology on qPCR-based methods and almost all were performed in a single laboratory. These previous studies (Fig 1 and Fig 2) included various sample sources and species and examined DNA extraction protocols that utilized phenol-chloroform, salting out, magnetic beads, and silica membranes from commercial kits with and without modifications. The majority of studies evaluated replicability by testing the correlation of TL measured in samples from which the DNA was extracted with different methods. Studies have not examined how different DNA extraction protocols affect assay precision nor have studies compared the resulting TL from each DNA extraction method with TL measured by different assay. All previous studies have reported TL measurement precision/reproducibility based on assay coefficient of variation (CV=S.D./mean value) rather than intra-class correlations (ICC). The ICC is a more appropriate measurement of assay precision for qPCR-based TL measurements because, for such measurements, the mean value used to calculate the CV is an assay-specific number without a true zero, and as such CVs are relatively arbitrary (19). Lastly, there has been wide variation in the methodologic approach to determine DNA quality, integrity, purity, and quantify dsDNA as potential indicators of the effect of different DNA extraction protocols on TL measurement. As such, while there is evidence that pre-analytic factors contribute to meaningful differences in the reproducibility of TL measurement and consequently also variability in the reported associations with TL, until now systematic investigation of the effects of DNA extraction, DNA integrity, and analytical methodology on the precision of TL measurement from blinded samples across multiple laboratories has been missing.

**Fig 1.**
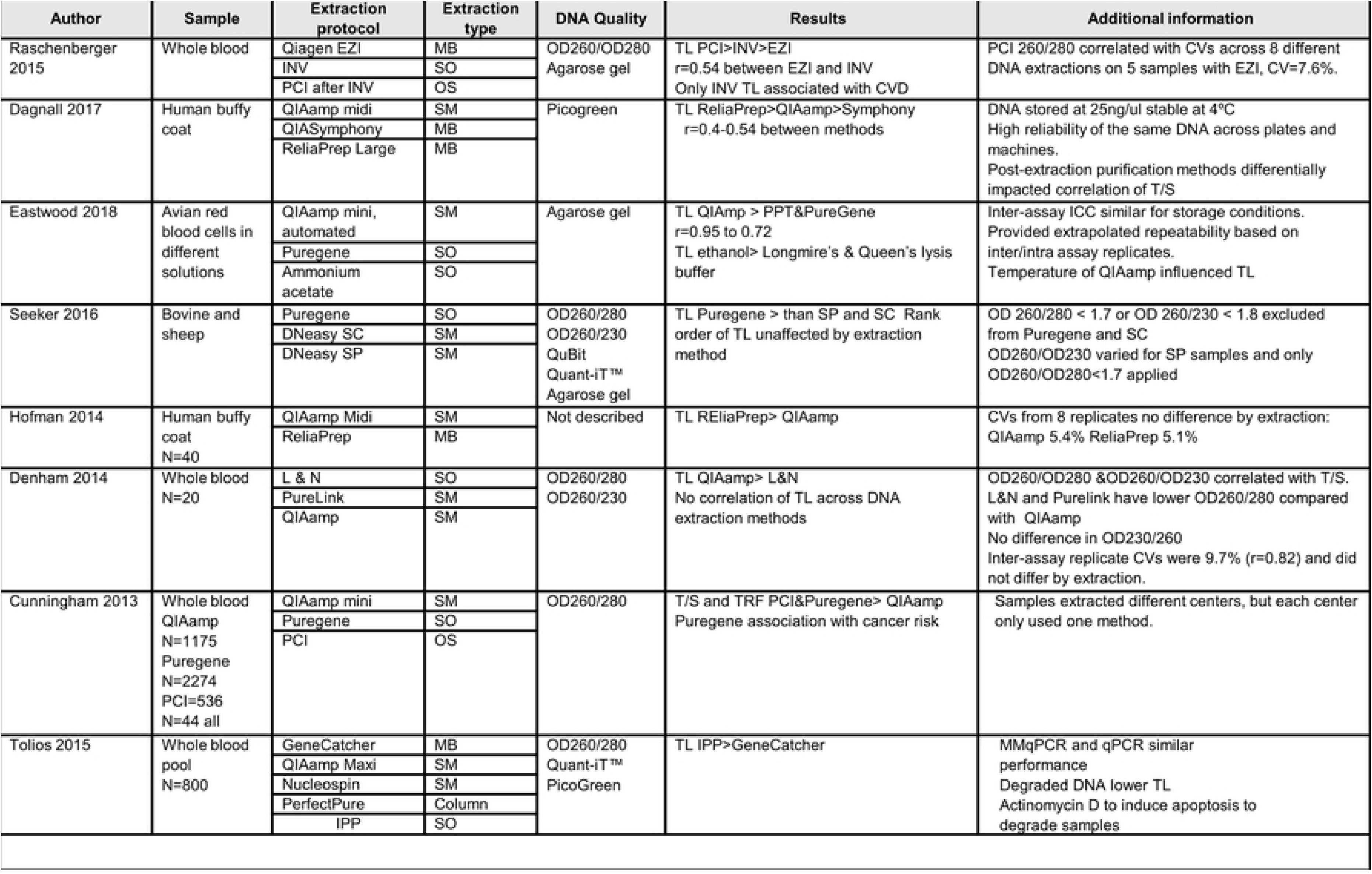
Summary of previous studies evaluating the impact of DNA extraction on telomere length measurement. Descriptions of biological sample source, DNA extraction protocols, quality control metrics and main findings. MB: magnetic beads; SM: silica membrane; SO: salting out; PCI: Phenol chloroform isoamyl alcohol; OS: organic solvent; INV: Invisorb® Spin Blood kit; IPP: Isopropanol precipitation; LN: Lahiri & Nurnberge

**Fig 2.**
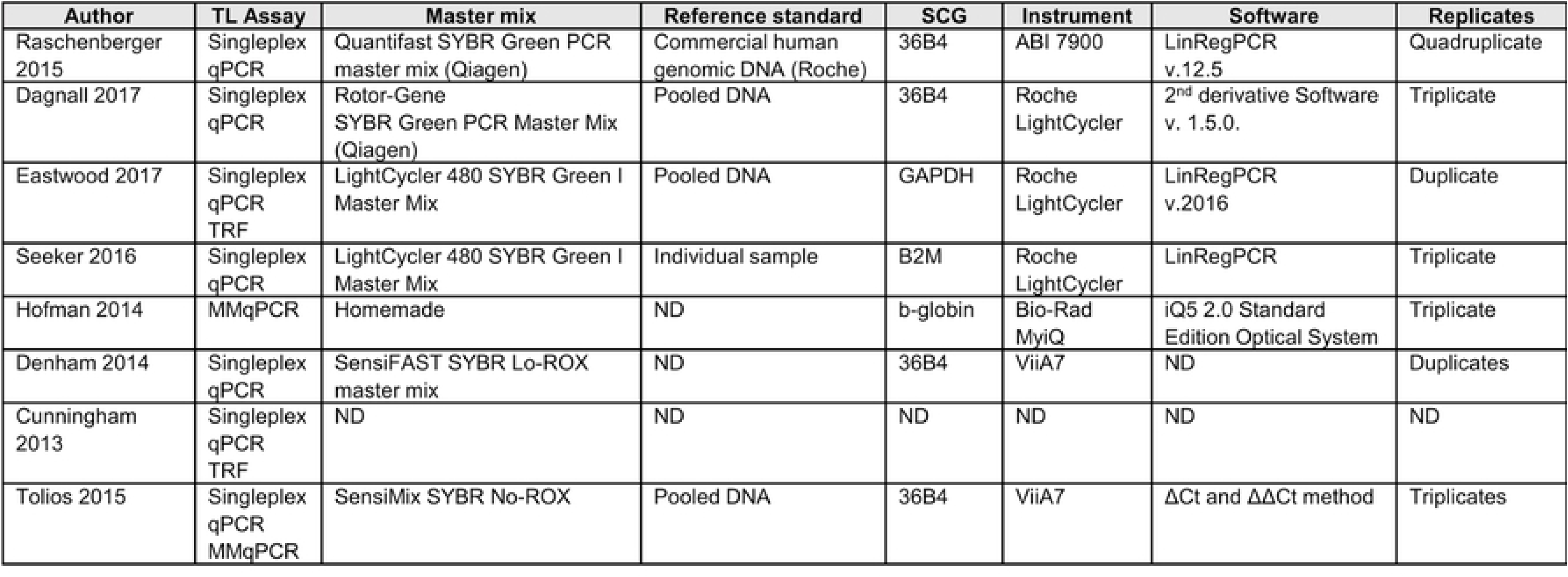
Summary of telomere length measurement methodology utilized in previous studies evaluating the impact of DNA extraction on telomere length measurement.

To address this critical gap, we selected three principal DNA extraction methods utilized frequently in population-based TL studies-salting out/precipitation (SO), silica membrane (SM), and magnetic beads (MB). Due to the infrequency of use in current telomere population-based studies we did not include a phenol-chloroform extraction method. We then selected commercially available kits that reflected the most often utilized for each method reasoning that this would provide more generalizable guidance for investigators. We selected whole blood as the source for DNA due to the predominance of this biological sample source in population-based studies. Anonymized donor whole blood samples that were from diverse racial/ethnic backgrounds and a wide age range of the participant were selected. All samples were blinded by an independent laboratory with statistical analyses performed by an independent statistician separate from all participating laboratories to ensure impartiality. When designing this study, we adopted an approach that reflected “real work” scenario, with each participating lab applying DNA extraction protocols routinely used in their lab to prevent effects related to the implementation of a novel extraction method. Examination of precision occurred at the level of the lab-protocol combination, as such we were able to specifically test the precision of TL measurement as a function of DNA extraction protocols, while accounting for the variability in TL assay measurement across labs. Using this approach, this study sought to do the following: (1) Systematically review the existing literature on the influence of pre-analytic factors on TL measurement; (2) Determine the precision qPCR-based TL measurement defined as a) the ICC of repeated TL measurement, performed on different days, but utilizing the same DNA sample; b) the ICC of repeated TL measurement from duplicate DNA samples extracted on two separate occasions from the same original blood sample; (3) Evaluate and compare the effect of DNA extraction methods on the precision of qPCR-based TL measurements; (4) Evaluate how commonly utilized metrics of DNA quality, purity and integrity influence qPCR-TL measurement precision.

## Methods

### Sample collection

Whole blood was obtained from a convenience sample of 50 individuals who ranged in age from 27 to 84 years, with a mean age of 57.7 years (S.D. = 12.6). The majority (82%) of the donors were male and the sample represented a diverse ethnic background (S1 Fig). Blood was collected into 6 tubes of 6 ml EDTA tubes (purple top, K2-EDTA, spray-coated), combined and aliquoted into multiple sets of 0.5 ml or 2 ml total within 4 hours of blood draw and stored at - 80°C until shipped to the independent U24 laboratory (SD).

### Blinding

Steps were implemented into the study design to ensure complete blinding of all samples at multiple levels (Fig 3). Each donor was assigned a random ID and the data linking the original subject demographics to this first blinding of IDs was sent separately to the statistician (SV) to ensure that no other individual was able to break the blinding. These aliquots were subsequently shipped on dry ice with continuous temperature monitoring (Onset InTemp Loggers, Bourne, MA) to the independent U24 laboratory (SD). Upon arrival to the U24 laboratory, each aliquot was assigned a new random ID by the data manager (CF) and re-labeled with no additional freeze thaw cycles. As with the previous blinding, the data linking the original blinded sample IDs to the secondary sample IDs was sent directly to the statistician to ensure no other individual was able to break the subject blinding.

**Fig 3.**
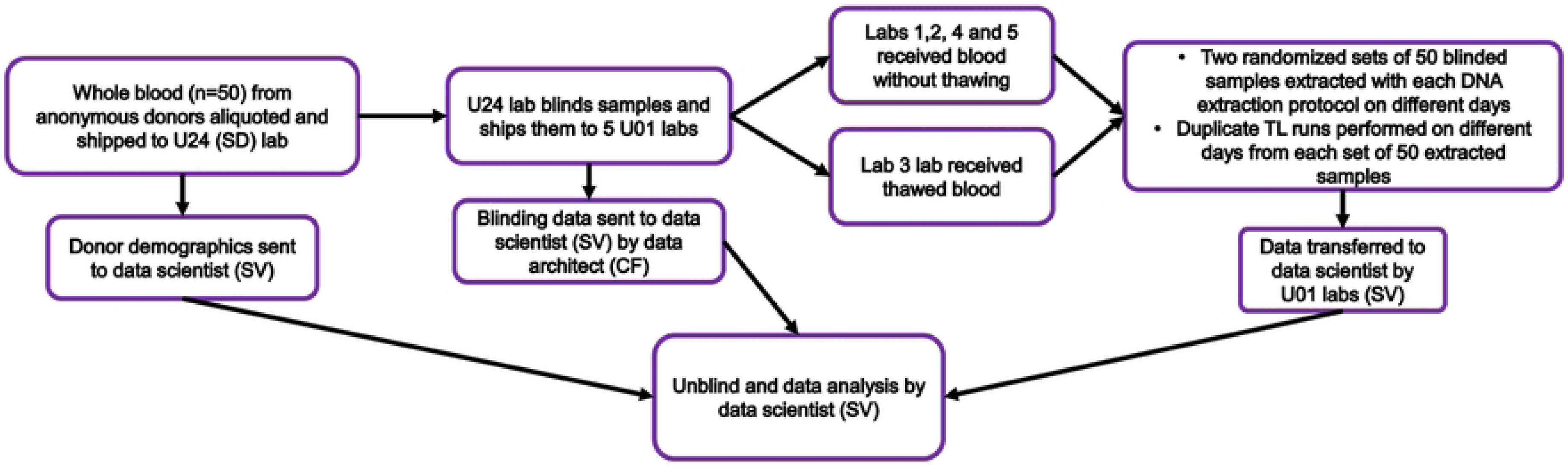
Study workflow.

### Sample distribution

The double-blinded aliquots were shipped on dry ice with continuous temperature monitoring to the 5 participating laboratories. Due to shipping difficulties, samples sent to one of the laboratories (Lab 3) were thawed upon arrival and experienced a range in temperatures, with a maximum of 17.9°C. After consideration of the “real world” study design, it was decided to proceed with the DNA extraction and subsequent TL assays. These samples were extracted with the QIAamp DNA Mini Kit and Gentra Puregene kit (QIAGEN). One additional lab (Lab 1) experienced temperature fluctuation in one of two separate shipments with a maximum temperature of −36°C, but no sample thawing. All other shipments maintained temperatures below −60°C during shipment. In total there were 10 duplicate sets of blood utilized to conduct the experiments described below. The overall study workflow is illustrated in Fig 3.

### DNA extraction protocols and laboratory allocation

Six different DNA extraction protocols reflecting the three most common DNA extraction types-magnetic beads, silica membrane-based columns and salting out/precipitation methods-were selected. Each DNA extraction protocol was conducted in at least two different laboratories and extractions were done according to the manufacturer’s instructions. Table 1 outlines the DNA extraction protocols, types and QC methods by laboratory. Each set of 50 samples was extracted separately on different days and in many cases by different technicians. Each set of 50 samples was analyzed on two distinct days for a total of 100 TL measures for each set of 50 samples. In the laboratory performing TRF the same technician performs all DNA extractions and performs all TRF assays for each individual cohort or study as part of the laboratories’ standard procedures.

**Table 1.** DNA extraction method, kit, and quality control metrics provided by participating laboratories. Laboratories 1-4 measured TL using qPCR, and laboratory 5 used Southern blot.

| Extraction Method | DNA Extraction Protocol | Lab | DNA QC Methods |
| --- | --- | --- | --- |
| **Magnetic bead (MB)** | Agencourt GenFind V2 (AG) | 1 | NanoDrop 2000C |
|  | Mag-Bind Blood & Tissue (KF) | 2 | Quant-iT PicoGreen dsDNA |
|  | QIAasymphony DSP DNA Midi (QS) | 2 | Quant-iT PicoGreen dsDNA |
|  | QIAasymphony DNA Mini (QS) | 4 | Nanodrop, TapeStation DIN |
| **Salting out/**<br>**Precipitation (SO)** | Gentra Puregene (PG) | 1 | NanoDrop 2000C |
|  | Gentra Puregene (PG) | 3 | NanoDrop |
|  | Gentra Puregene (PG) | 4 | Nanodrop, TapeStation DIN |
|  | Gentra Puregene (PG) | 5 | Nanodrop, agarose gel |
| **Silica membrane (SM)** | QIAamp DNA Blood Midi (QD) | 1 | NanoDrop 2000C |
|  | QIAamp DNA Blood Mini (QN) | 1 | NanoDrop 2000C |
|  | QIAamp DNA Blood Mini (QN) | 3 | NanoDrop |
|  | QIAamp DNA Blood Mini (QN) | 4 | Nanodrop, TapeStation DIN |

### Telomere length analyses

To best understand the specific impact of DNA extraction on assay precision across DNA extractions within a laboratory, TL was assayed using the specific standardized TL assay in that laboratory. Four laboratories performed singleplex qPCR and qPCR results are expressed as the ratio of telomere signal (T) vs. a single-copy gene (S), i.e. T/S ratio. One laboratory utilized a Southern blot, Telomere Restriction Fragment length (TRF) analyses and TRF results are expressed in base pairs. Key features of analytical factors within the qPCR lab protocols are listed in Table 2. Full protocols for all laboratories are available as supplemental material and listed by the corresponding laboratory number in figures and tables. (S1 – S5 protocols)

**Table 2.** Summary of telomere length methodology utilized by participating laboratories. SCG: single-copy gene. *Restriction enzyme reported for TRF.

| Lab | TL Assay | Master Mix | Reference Standard | SCG* | Instrument | Software | Replicates |
| --- | --- | --- | --- | --- | --- | --- | --- |
| 1 | Singleplex qPCR | Homemade | Commercial DNA (Sigma-Aldrich) | HBG | Roche LightCycler 480 | LightCycler v1.5.1 | Triplicate |
| 2 | Singleplex qPCR | SYBR Green PCR Master Mix (QIAGEN) | Pooled DNA | 36B4 | Roche LightCycler 480 | LightCycler v1.5.0 | Triplicate |
| 3 | Singleplex qPCR | Homemade | Pooled DNA | 36B4 | Applied Biosystems | QuantStudio | Triplicate |
| 4 | Singleplex qPCR | SYBR Green PCR Master Mix (QIAGEN) | oligomers (IDT) | IFNB1 | Rotor-Gene Q | Rotor-Gene Q v2.1.0 | Triplicate |
| 5 | TRF | N/A | Commercial DNA ladder (Invitrogen) | *HinfI*, *RsaI* * | N/A | ImageQuant | Duplicate |

### DNA quality measurements

To assess specific factors related to DNA quality, purity and integrity, several analyses were conducted. All laboratories sent their “routine” QC metrics to the TRN statistician (SV). Lab 5 evaluated DNA integrity via agarose gel electrophoresis only. All data was compiled in relation to Nanodrop and Picogreen/Qubit assessment of DNA concentration and purity. The Nanodrop provides a measure of nucleic acid concentration as well as two ratios, the OD260/230 and the OD260/280 that are derived from the absorbance at these wave lengths with nucleic acid absorption (both DNA and RNA) at 260nm. In general, the OD260/230 ratio is considered an indicator of sample contamination with substances that absorb at 230nm including salts, EDTA, and phenol. As such a low 260/230 ratio indicates impurities in the sample and a ratio greater than 2.0 is generally considered acceptable. Protein absorption occurs at a wavelength of 280nm, as such a low OD260/280 ratio is indicative of protein contamination in the sample and an OD of 1.8 generally thought to indicate pure DNA and an OD of 2.0 to indicate pure RNA. As protein, EDTA, and other impurities can impact PCR efficiencies, primer binding, and enzyme activity, variations in these impurities between samples, or replicates, can potentially affect measurement precision. To evaluate this, we explored the relation between these commonly used indicators and the ICC calculated over repeated extractions.

In addition to the reporting of the standard QC data obtained at each laboratory, after the completion of TL analyses, the remaining DNA from all participating laboratories was returned to the U24 laboratory for the assessment of genomic DNA integrity using the Agilent TapeStation. Samples were shipped overnight, on dry ice, with continuous thermal monitoring. No temperature deviations in any of the shipments were observed. The Agilent TapeStation generates a numerical assessment of global genomic integrity, the DNA integrity number (DIN), based on an algorithm of over 7000 genomic DNA samples (12). DIN was measured on the exact same 6 individuals from each laboratory and each duplicate DNA extraction for a total of 6 samples with 12 sets of duplicate DNA equaling 144 independent DIN measurements. Measuring six samples was deemed sufficient based on preliminary DIN data showing variation within batched of extracted samples to be very small relative to variation between batches of extracted samples.

### Data collection, processing and data flow

All data derived from the TL assays were integrated following a multi-level procedure governed by pre-specified data validation workflows. Briefly, each participating laboratory structured its independently generated TL data into an import template provided by the data architect (CM). This template had pre-configured data attributes, input masks, and in-field validation queries to ensure the completeness and consistency of the collected data. Upon finalizing data collection, each laboratory transmitted a locked import template to the data manager following a data transfer workflow via a secured digital channel. All received data were automatically integrated and indexed into a secured cloud-based data warehouse. Multi-level, post-hoc curation processes were automatically triggered to ensure logical and physical data integrity. Any compromised data points were flagged as part of the process, and the data architect (CF) conducted detailed verification of each value with the corresponding source laboratory prior to locking the database. After finalizing the data validation process, a unified long-format annotated data set was generated and transmitted to the statistician (SV) for analysis (Fig 4).

**Fig 4.**
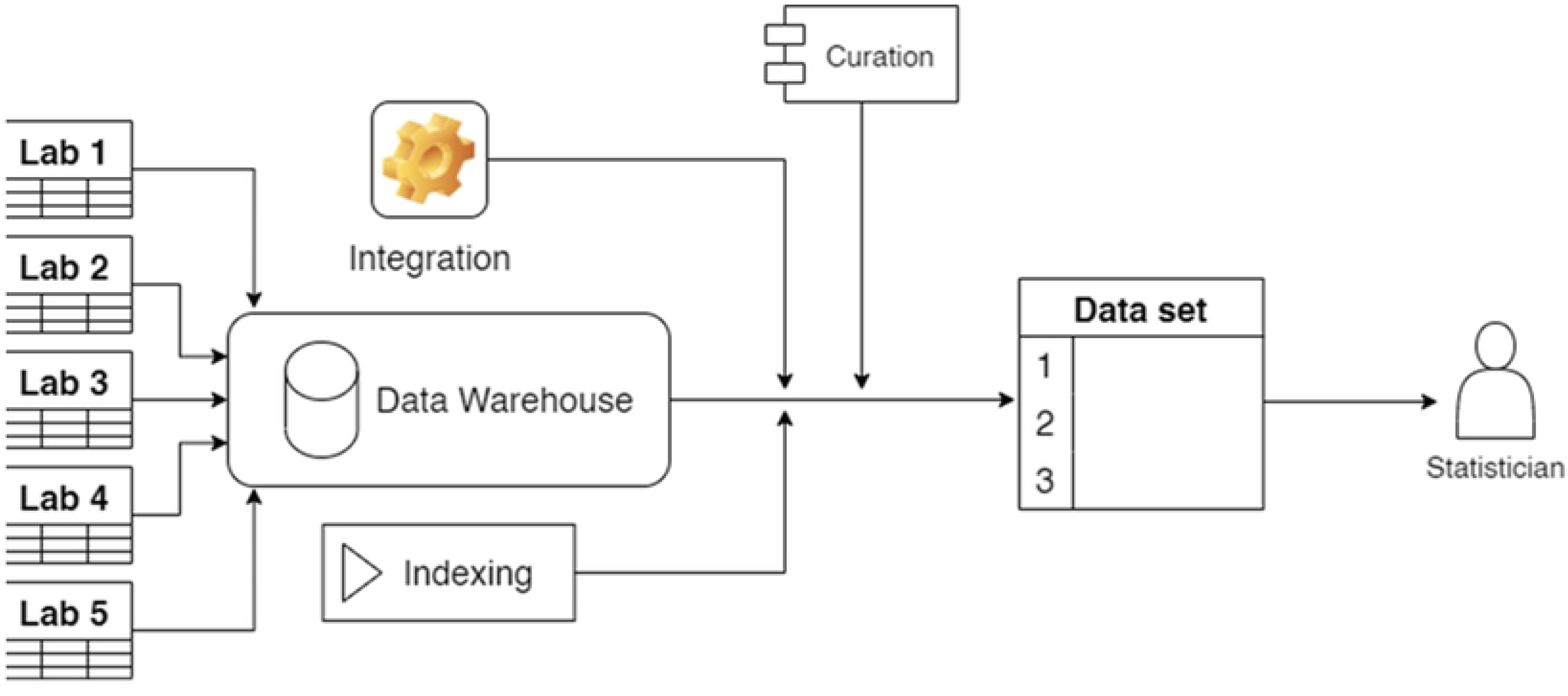
Data flow. Data flow from labs via the data warehouse to the statistician

### Statistical analyses

Prior to analysis, T/S ratios were Z-transformed (mean = 0, S.D. = 1) per measurement set (n=50) (i.e. for each lab / protocol / extraction / qPCR run combination), to eliminate assay specific effects on mean and variance of T/S ratios that would affect the ICC estimates.

ICC estimates were calculated using the R-package rptR (13). ICC estimates vary between zero and 1, with zero indicating no correlation at all (zero precision) between different measurements, and 1 indicating a perfect correlation (maximum precision) between different measurements (see Fig 5 for illustration of data yielding different ICC estimates). Since age and sex effects are usually known in telomere studies of human subjects, and ignoring age and sex on the T/S ratio would inflate the ICC estimates due to the increase in variance, all ICC estimates are from models that included age and sex as fixed effects (12). When estimating the ICC over repeated measurements of the same extraction of a sample, the different extractions of each sample were treated as if they were from different samples as this was a straightforward way to obtain a single estimate over all data per lab / protocol combination.

**Fig 5.**
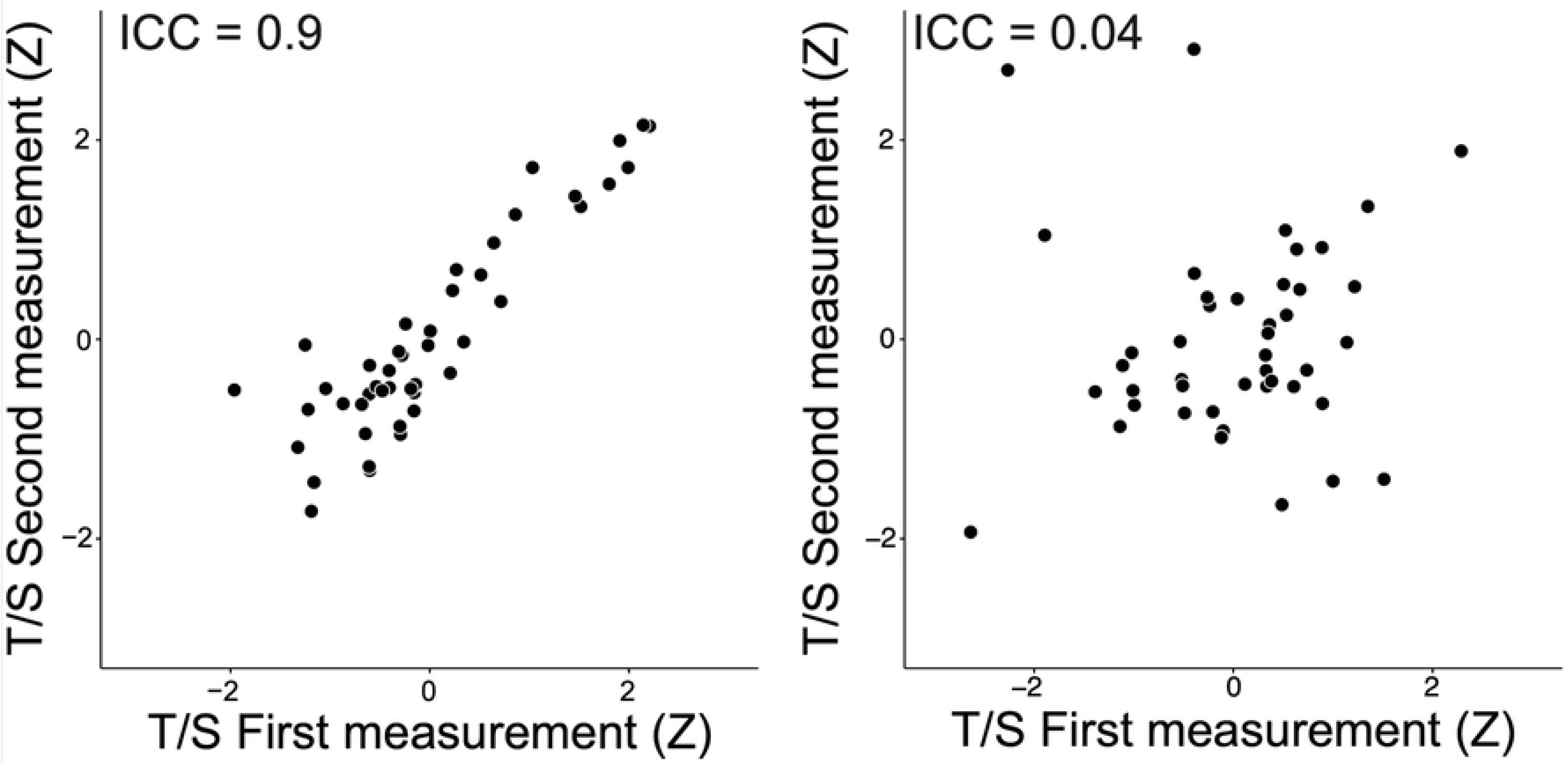
Scatterplots of example z-scored T/S ratios from this study. A. a sample with high (ICC 0.9) B. a sample with low ICC (ICC 0.04) estimate. Note that in this study the ICC estimates all refer to ‘correlations’ between two measurements, and thus superficially resemble Pearson correlations, but ICC estimates can equally be calculated over more than two measurements per sample.

Pearson correlations between T/S ratios and TRF were calculated for each lab / protocol / extraction / qPCR run combination (n=50 per correlation) from a model including the T/S ratio only, i.e. without other variables.

Statistical comparisons between ICC estimates were performed using a meta-analysis approach, to account for the uncertainty in the ICC-estimates (ignoring this uncertainty would underestimate confidence intervals and p-values). This analysis involved testing for effects of laboratory identity and protocol effects, including these as factors, and testing for effects of quality control measurements of the DNA-samples, including these as co-variates (i.e. a meta-regression approach). Due to the modest sample size, independent variables were tested one at the time. ICC estimates were Fisher-z transformed and weights were based on the inverse of the ICC variance. Meta-analyses were carried out using the rma-mv function of the R-package metaphor, with QM as test statistic of the Wald-type test (with associated d.f. noted as subscript) of the model coefficients (14).

## Results

### Telomere length measurement precision of same DNA aliquot run on different days

We first examined the precision of TL measurement of the same DNA sample run on different days (Table 3). This analysis showed that ICCs of qPCR assays varied widely (range 0.43 to 0.94) and were lower compared to the ICC of Southern blot (0.99) (Fig 6A). The analysis further revealed large variation between labs (qPCR only: QM_3_=105.6, P<0.0001), relative to the variation between DNA extraction protocols within the same lab. There were no consistent effects of extraction kit (QM_5_ = 4.10, P=0.53) and type (QM_2_ = 0.76, P=0.68) on TL measurement precision of duplicate measurements utilizing a single DNA sample performed on different days.

**Fig 6.**
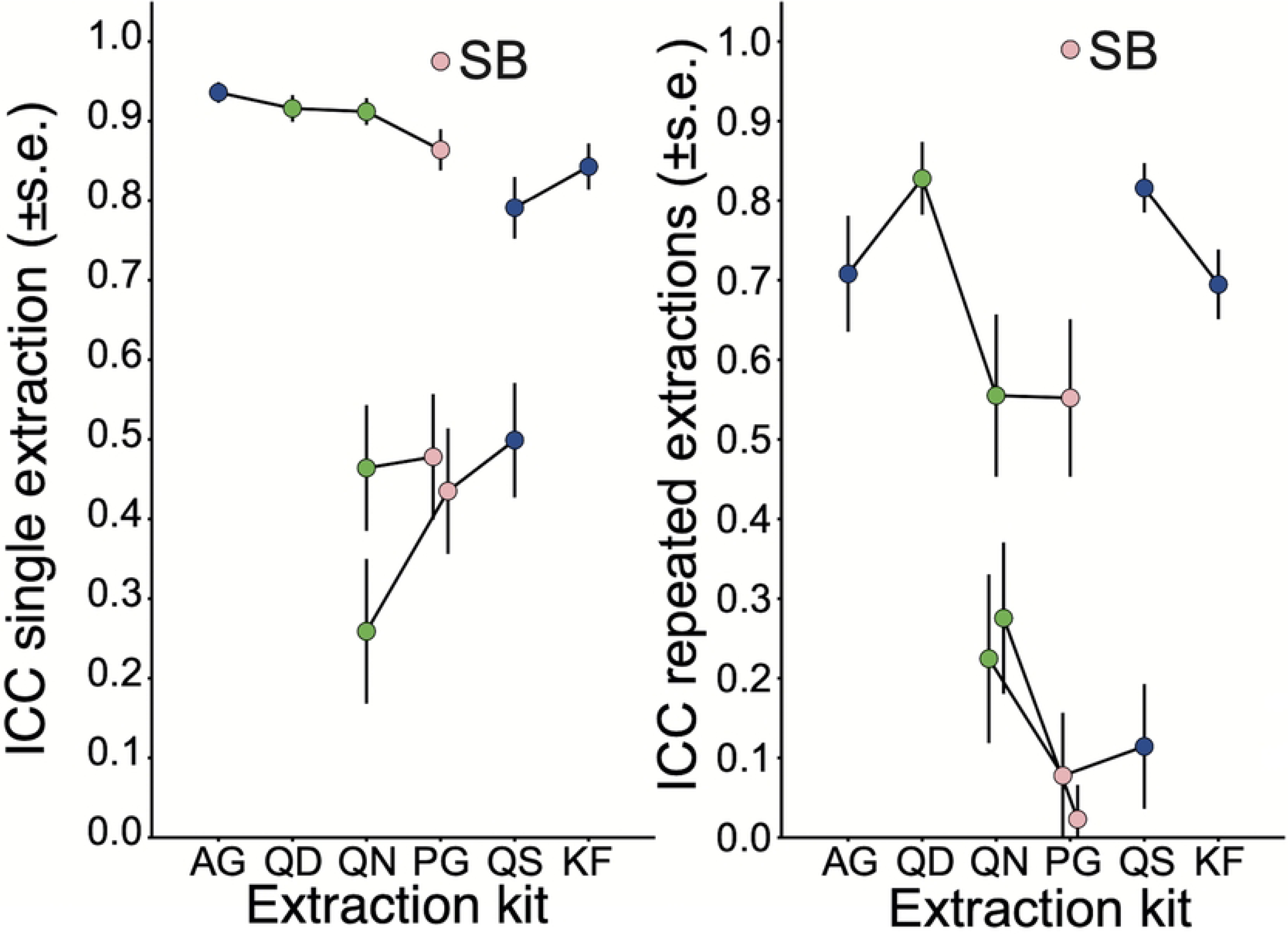
Telomere measurement precision as indicated by the Intraclass Correlation Coefficient. A. Intraclass correlation coefficient (ICC, ±s.e.) calculated over measurements of a single extraction of each sample B. ICC calculated over measurements of two extractions of each sample. Each data point represents one laboratory / extraction protocol combination – all qPCR based except for the data point labelled SB (Southern blot). Data points connected by a line are from the same laboratory. Abscissa shows the different extraction kits (AG: Agencourt; QD: QIAamp Midi; QN: QIAamp Mini; PG: PureGene; QS: QIAsymphony; KF: KingFisher). Data point colors indicates extraction principle (blue: magnetic beads; green: silicone membrane; pink: salting out). Where there appears to be no s.e. this was smaller than the marker.

**Table 3.** Measurement precision of TL measurements by DNA extraction method and protocol. Precision estimates (ICC ± standard error) were calculated over repeated measurements of one extraction per sample (i.e. same DNA) and over measurements on repeated DNA extractions of each sample. Also shown is the correlation between qPCR-based TL and TRF.

| Lab | Extraction Method | Extraction Protocol | ICC – same DNA extraction | ICC – repeated DNA extractions | R<sup>2</sup> - correlation with TRF |
| --- | --- | --- | --- | --- | --- |
| 1 | MB | AG | 0.936±0.013 | 0.708±0.073 | 0.700 |
| 2 | MB | KF | 0.843±0.029 | 0.695±0.044 | 0.773 |
| 1 | SO | PG | 0.864±0.026 | 0.552±0.099 | 0.501 |
| 4 | SO | PG | 0.435±0.079 | 0.078±0.079 | 0.166 |
| 3 | SO | PG | 0.478±0.079 | 0.023±0.043 | 0.013 |
| 5 (TRF) | SO | PG | 0.975±0.005 | 0.99 | - |
| 1 | SM | QD | 0.916±0.017 | 0.828±0.046 | 0.760 |
| 1 | SM | QN | 0.912±0.017 | 0.555±0.102 | 0.655 |
| 4 | SM | QN | 0.259±0.091 | 0.225±0.106 | 0.288 |
| 3 | SM | QN | 0.464±0.079 | 0.276±0.095 | 0.231 |
| 4 | MB | QS | 0.499±0.072 | 0.114±0.079 | 0.214 |
| 2 | MB | QS | 0.791±0.039 | 0.816±0.031 | 0.782 |

### Telomere length measurement precision of repeat DNA extraction from the same blood sample on different days

We reasoned that assessment of the impact of DNA extraction methods on TL measurement precision needs to encompass the entire process, which starts from an aliquot of whole blood. To this end, each set of whole blood, which included 2 randomly blinded aliquots from each of the 50 donors, was extracted twice with the same DNA extraction kit on different days and assayed independently. Fig 6B shows color coded plots of these analyses. The ICCs of qPCR assays were lower than the ICC of SB (Fig 6B). The ICCs calculated over measurements on duplicate extractions (Table 3) were on average 0.23 lower than the ICCs calculated over duplicate measurements of a single extraction per sample (qPCR results only: z=4.3, P<0.001; Fig 7). Similar to the findings with the same DNA samples, the precision varied strongly between labs (QM_3_=29.3, P<0.0001), even with the same DNA extraction protocol. There were no consistent effects of extraction protocol (QM_5_=5.84, P=0.32) or method type (QM_2_=2.63, P=0.27) on TL measurement precision calculated over measurements on duplicate DNA extractions per sample when the combined data from all four labs were considered.

**Fig 7.**
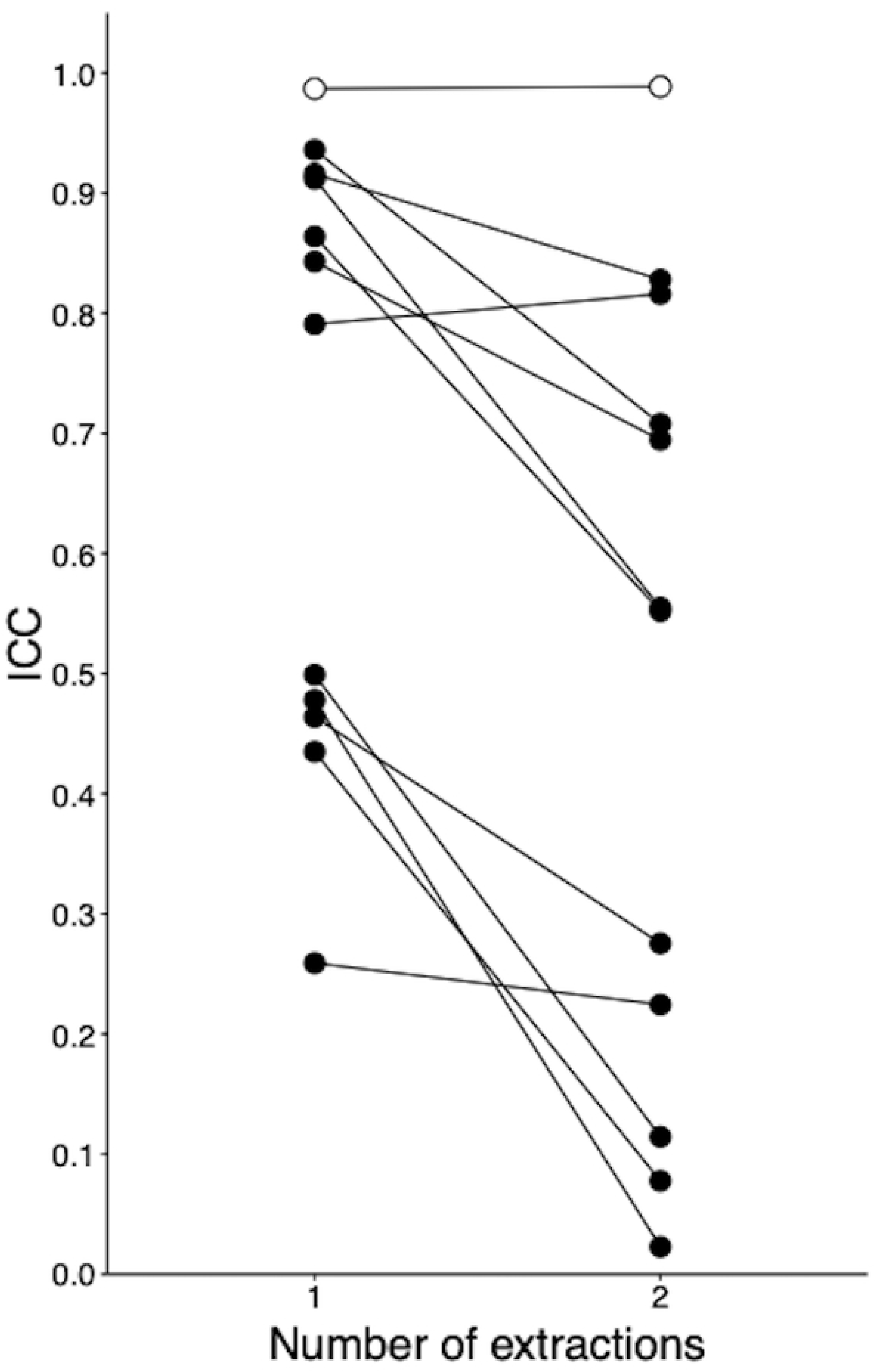
Extraction effect on measurement precision. Extraction effect indicated by the ICC calculated over measurements of a single extraction versus measurements of two extractions (number of extractions). Lines connect the ICC estimates for each laboratory / extraction protocol combination. All are qPCR based except for the data points labelled SB (Southern blot). See Fig 6 for the standard errors.

Within each protocol, the three ICC-estimates were lower for lab 4. This lab-to-lab difference is likely due to the multiple analytical factors associated with each lab’s qPCR protocol (Table 2) and will be the focus of future investigations. Despite all of these other sources of variation, within a lab (labs 1, 3, 4), the Puregene kit (salting out type) had the lowest ICC values for all three labs. No consistent pattern was observed for the silica membrane and magnetic beads types. However, magnetic beads (AgenCourt for lab 1, QIAsymphony and KingFisher for lab 2) appear to have overall higher ICCs between repeat extractions.

### Cross methodology test of implications of ICC variability

To evaluate the relevance of the wide variability in ICC, and to better understand the implications of the differences in ICC between the same DNA assayed on different occasions and extraction of the same blood sample, we utilized the relation between qPCR-based TL measurement and TL determined by TRF. Specifically, we calculated the correlation between the TL estimates obtained using qPCR and TRF measured on independent extractions of the same samples for each lab / protocol combination, as an additional measure of the relevance of the variability in ICC for the qPCR-based assays (Table 3), assuming that a higher correlation between T/S ratio and TRF indicates higher measurement precision. The strength of these correlations varied widely, from R^2^=0.01 to 0.78. The ICCs calculated over TL measurements of duplicate extractions predicted these R^2^ well (R^2^=0.95), while the ICCs calculated over repeated TL measurements on single extractions of each sample performed less well in this respect (R^2^=0.52; Fig 8). The lack of overlap between the 95% confidence intervals of these estimates (not shown) are indicative of a significant difference. Similar results were obtained using the association between T/S ratio as dependent variable and age and sex as independent variables (S2 Fig).

**Fig 8.**
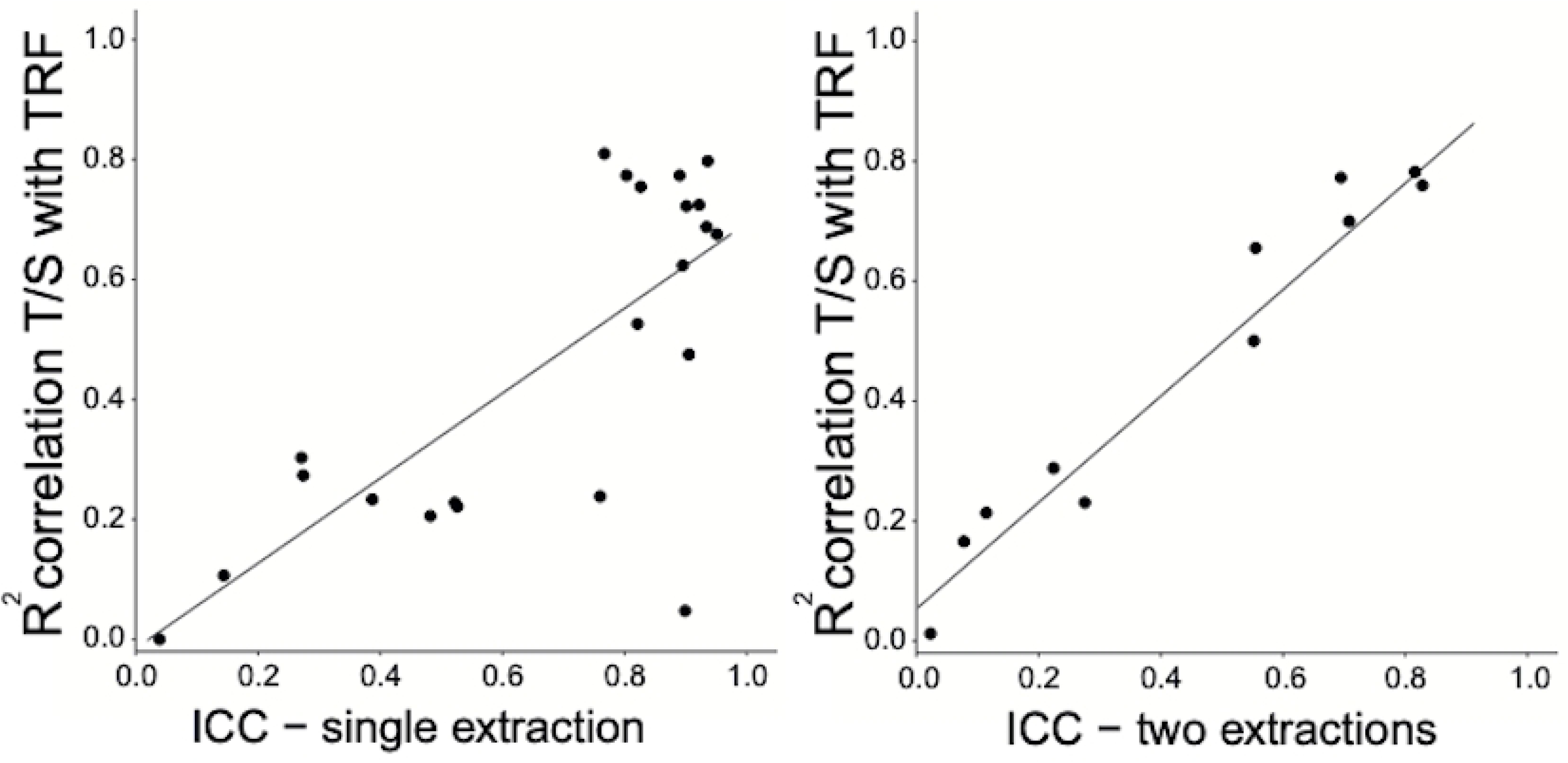
Comparison of precision metrics. The Y-axes show the R^2^ of the associations between T/S ratios and TRF on the same samples, and the abscissa shows two metrics of measurement precision (Intraclass Correlation Coefficient, ICC). Each data point represents an R^2^ value calculated over the 50 samples in each set. A. ICC calculated over measurements of a single extraction of each sample, and 8B. ICC calculated over measurements of two extractions of each sample. Each laboratory / extraction protocol combination is represented by two data points in panel 8A (one for each of the two extractions) and one data point in panel 8B. The correlation in panel 8B is significantly stronger than the correlation in panel 8A.

### Sample quality effects on the ICC

Given the wide variation in ICCs observed for qPCR-based TL measurements, and the recognition that DNA quality and purity may be influential on qPCR-based biological assays that rely on accurate primer binding, polymerase activity, and PCR efficiency, we explored how the quality of the extracted DNA, assessed using the OD260/230, OD260/280, and the DNA integrity number (DIN) obtained via a TapeStation bioanalyzer contributed to the variation in the ICC between labs and protocols (S3 Fig). The DINs of repeated extractions of the same blood sample were highly correlated (S4 Fig) suggesting that DIN is an indicator of the sample itself.

Using a meta-regression approach, we tested for associations between three DNA sample quality metrics, DIN, OD260/230 ratio and OD260/280 ratio on the one hand, and the ICC on the other hand (Fig 9). All three DNA sample quality metrics were positively associated with the ICC (Table 4). This association reached significance for the OD260/230 ratio only (lower OD260/OD230 indicating contamination with organic compounds), while associations between ICC and the DIN (DNA Integrity Number), and the OD260/280 ratio (lower ratio indicating protein contamination) were trends (Table 4). However, the associations were similar in strength, with the lowest and highest p-value only 0.062 apart. On the basis of the current analysis, there is insufficient support to weigh one quality metric over any other assessment of DNA quality and integrity when evaluating the effect on TL measurement precision. We conclude that all three quality metrics are somewhat predictive of the ICC of qPCR-based TL measurements. However, we note that high scores on each of these metrics, suggestive of better integrity or DNA quality, can still result in low ICC values (Fig 9).

**Fig 9.**
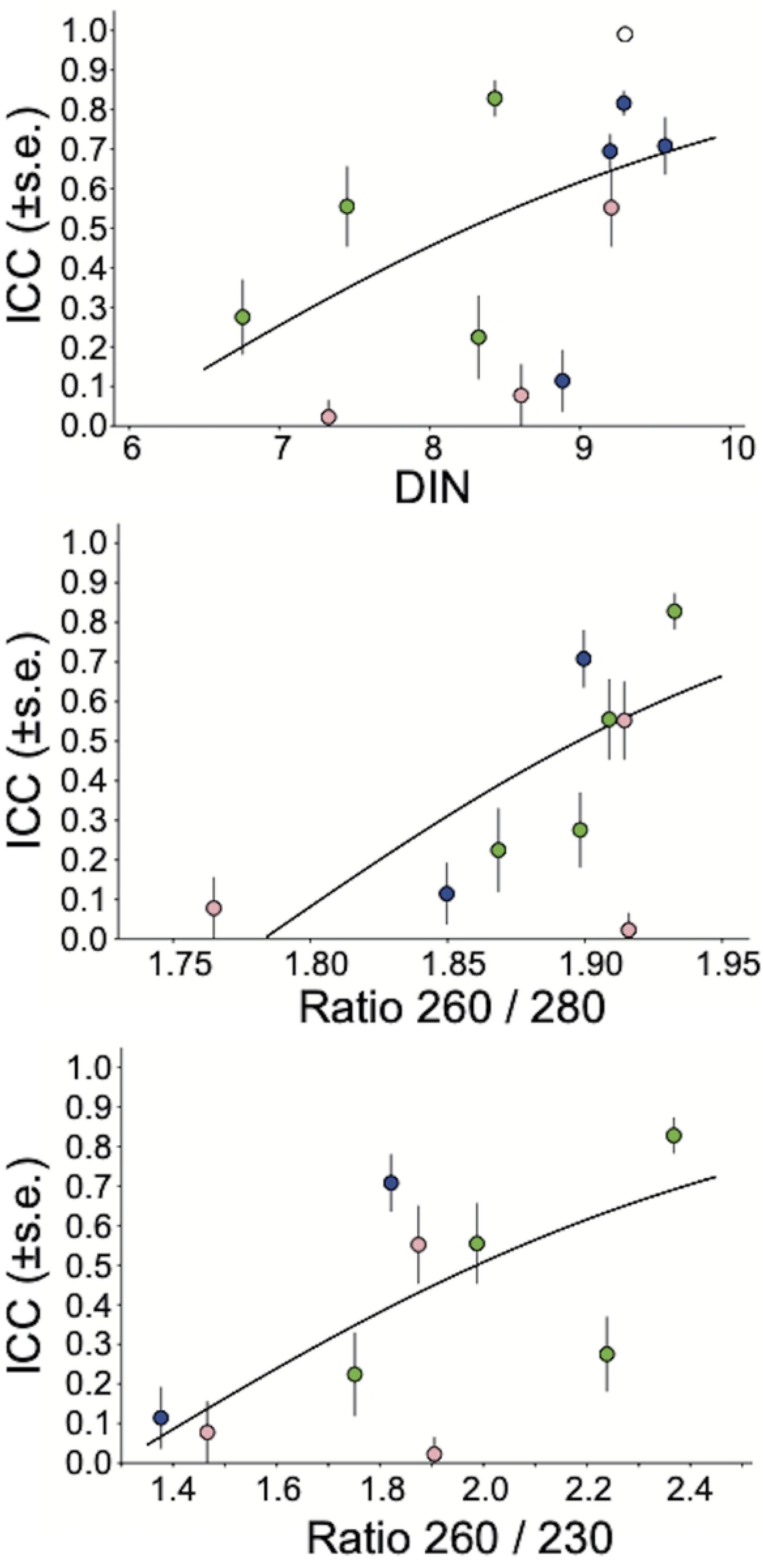
Measurement precision in relation to sample quality control metrics. A. DNA Integrity Number (DIN), B. Contamination with proteins (260 / 280 ratio), C. Contamination with organic compounds (260 / 230 ratio). Regression line shown was back-transformed (z to r) from equation in Table 4.

**Table 4.** Effect of DNA sample quality on qPCR-based precision meta-regression. DNA sample quality control metrics as univariate predictors of the ICC (Fischer-z-transformed) of qPCR-based telomere measurements calculated over different extractions. n=11 for DIN, and n=9 for others reflecting one per lab / DNA extraction protocol combination. Z- and P-values refer to the coefficient of the quality metric.

| Metric | Intercept ± s.e. | Coefficient ± s.e. | Z | p |
| --- | --- | --- | --- | --- |
| DIN | -1.35±1.17 | 0.23±0.14 | 1.16 | 0.098 |
| OD260/230 ratio | -1.02±0.72 | 0.79±0.38 | 2.10 | **0.036\*** |
| OD260/280 ratio | -8.54±4.66 | 4.79±2.47 | 1.94 | 0.053 |

DNA sample quality metrics OD260/230 ratio and OD260/280 ratio varied also within the batches of 50 samples, and we tested the extent to which this variation was associated with the ICC. To this end, we divided the data from each lab / protocol / extraction combination where the QC data were available in two sets (n=25 each) for each quality metric, with the lowest 25 QC values in one group and the highest 25 QC values in the other group. We then calculated the ICC for these groups separately, using the repeated TL measurements on each extraction (using the ICC calculated over multiple extractions was problematic due to each sample having two different QC values, one for each extraction, but, as illustrated in Fig 7, there was a strong correlation between ICC estimates of single and multiple extractions). Visual inspection of the data shows that within batch variation in the OD260/280 ratio was not associated with the ICC (panel A in S5 Fig). In contrast, in nine out of ten batches the ICC of the set with the higher OD260/230 ratio was higher (panel B in S5 Fig), indicating a strong within batch association between the OD260/230 ratio and the ICC.

On the basis of the current analysis, there is insufficient support to weigh one quality metric over any other assessment of DNA quality and integrity when evaluating the effect on TL measurement precision. We conclude that all three quality metrics are somewhat predictive of the ICC of qPCR-based TL measurements, with the added observation that the OD260/230 ratio also predicts precision within batches, while the OD260/280 ratio does not share this property. However, we note that high scores on each of these metrics, suggestive of better integrity or DNA quality, can still result in low ICC values (Fig 9).

## Discussion

We systematically evaluated the effect of repeated DNA extraction on the precision, e.g. the ICC of repeated TL measurement of the same sample, of qPCR-based TL assays across laboratories and across DNA extraction protocols. The fully blinded multi-lab study design permitted the evaluation of both DNA extraction protocol and lab effects on measurement precision. Overall, the precision of qPCR-based methods was significantly lower than Southern blot, consistent with earlier findings (15, 16). Our findings that both DNA extraction protocol and lab-specific methodology influence precision are consistent with the previous literature and have several direct implications for current and future studies. First, studies expecting to include samples extracted with different DNA extraction protocols should proceed with significant caution, and if possible, avoided. The inclusion of TL measured on DNA utilizing different DNA extraction protocols, and for that matter measured in different labs, is predicted to impact the overall ICC of the study. Investigators should therefore evaluate this in light of the known effect of lower ICC on statistical power (1). Second, multi-site large scale studies should utilize uniform DNA extraction protocols when feasible. Additionally studies should maintain and report data on sample processing and storage, DNA extraction protocol, reagent batches, DNA storage, and specific machines utilized to generate TL data as the recent UK biobank study demonstrated clear effects of multiple pre-analytic factors on TL measurement (17). The TRN website also contains various resources to facilitate investigators as they evaluate different study designs including a sample storage and collection checklist (https://trn.tulane.edu/resources/study-design-analysis/) (2, 18–20). Third, when single DNA extraction protocols are not feasible, integration of protocol type into analysis, and careful evaluation for systematic differences should occur prior to hypothesis testing with TL. Our results also have implications for studies examining TL across multiple tissues. Systematic testing of the effect of different DNA extraction protocols on TL precision measured from different tissue sources has not been done. However, our results are consistent with reported differences in TL and precision measured in saliva, peripheral blood mononuclear cells (PBMCs) and dried bloods spots (DBS) from the same individual, where salting out DNA extraction was utilized for saliva extraction but Purelink, a silica membrane-based kit, was utilized for the DBS and PBMCs (21). Studies seeking to determine the true correlation of TL measured from different tissue sources in the same individual will need to address variation induced by different DNA extraction protocols.

We extend existing literature by finding a significant effect of repeated DNA extraction from the same blood source on qPCR assay precision-defined as the ICC between duplicate independently extracted samples. More specifically, we find that the ICC calculated over measurements of different DNA extractions, from the same sample, were substantially, and significantly, lower than the ICC calculated over multiple TL measurements of the same DNA sample. This indicates that the extraction process itself affects the DNA in a way that affects the qPCR results, and hence measurements on a single extraction, however precise, are specific not only to the sample, but also to that specific extraction. The ICC calculated over multiple extractions was a better predictor of the strength of the correlation between T/S ratios and TL measured via TRF on the same samples, confirming the superiority of ICCs calculated over multiple extractions. We therefore conclude that the ICC estimated over repeated measurements of a single DNA extraction overestimates measurement precision, and that measurements on multiple DNA extractions, instead of repeated measurement of the same DNA, is required for an accurate estimate of qPCR TL assay precision. Eastwood et al (2018) reached a similar conclusion based on a single-lab study measuring telomere length with qPCR on avian DNA.(2) Obtaining an accurate estimate of measurement precision is critical because the statistical power of a study increases with increasing measurement precision. This relation between ICC and statistical power has an unusually strong impact for epidemiologic studies of TL, where effect sizes are typically small, in the order of 100-300 base pairs, relative to the typical variation in (age-adjusted) TL between individuals, generally around 3000 base pairs. As a consequence, epidemiological studies of variation in TL are expected to require large sample sizes, in the hundreds to thousands, in order to have sufficient power, even when TL measurements are relatively precise. (see Lindrose et al. 2021 for example power calculations) (1).

Our data indicate significant laboratory effects on measurement precision, while DNA extraction protocol did not have significant independent effects replicated across laboratories, making it challenging to provide clear, data-informed, statements about which DNA extraction protocol yields the greatest precision. Although the original study goal of providing recommendations relative to the specific DNA extraction protocol for qPCR TL is confounded by the between laboratory findings, due to consistent low ICCs from labs 1,3, and 4 (Table 3 & Fig 7), the use of the Puregene kit for singleplex TL estimates utilizing qPCR is not recommended. Eastwood et al (2018), when comparing Puregene with QIAmp, reached the same conclusion.(2) Retrospectively, a technical problem with a robotic pipetting instrument in one of the participating laboratories was discovered that likely contributed to the lower than expected ICCs. This “equipment failure” highlights the critical importance of assay platforms and the need for regular internal quality control for each set of analyses even in laboratories with decades of experience in the TL assay. Enhancement of the precision of qPCR-based TL measurements will require additional studies to tease out the unique and combined contributions of individual factors (e.g. master mix, single copy gene, etc.) in order to best provide guidance to achieve the highest consistent precision when utilizing qPCR-based assays for TL determination.

TL can be measured using a various methodologies, including both the broad categories of PCR-based measurements (e.g. singleplex, multiplex), non-qPCR based approaches that utilize fluorescence in situ hybridization (e.g Flow-FISH, Q-FISH), restriction digests (e.g (TRF), or combinations of approaches (e.g. TeSLa, STELLA) further contributing to variability in assay precision, in part due to the use of different reagents and approaches. Even within singleplex PCR-based assays there are significant protocol differences including the source of master mix (e.g. purchased compared to homemade), single-copy gene, source of reference standards, machine platform, number of replicates, etc. (Table 2). Similarly, southern blot-based assays also have a broad range methodologic differences between laboratories and across protocols, including the selection and number of restriction enzymes, analytic software utilized to determine signal intensity, and whether or not hybridization is done within a gel or transferred to a membrane before hybridization. Methods that combine PCR and blot-based approaches, such as TeSLA, exist and create further nuanced levels of methodological complexity wherein differences in precision may arise. While the scope of differences in the qPCR assay results with the same DNA extraction protocols were unexpected (Table 3 and Fig 7), it revealed the critical importance of assay platforms and the need for each lab, regardless of methodology or experience, to calculate, and report, ICCs as an indicator of assay precision for each cohort and study. Enhancement of the precision of qPCR-based TL measurements will require additional studies to tease out the unique and combined contributions of individual factors (e.g. master mix, single copy gene, etc.) in order to best provide guidance to achieve the highest consistent precision when utilizing qPCR-based assays for TL determination. Our conclusion that ICC calculated over measurements on repeated extractions is a better indication of measurement precision assumes that a high correlation between qPCR and TRF results indicates better qPCR assay precision.

Overall, the precision of qPCR-based methods was significantly lower than Southern blot, consistent with previous findings in nonhuman vertebrates.(16) This study, however, was not designed to evaluate how DNA extraction or methodology impacts TL measurement using Southern blot, and we are therefore unable to make conclusions about the precision of Southern blot across different DNA extraction protocols, laboratories, or other pre-analytical and analytical factors. Despite this limitation, our data indicate both significant laboratory and DNA extraction protocol effects, creating analytic challenges for clear statements about the DNA extraction type that yields the greatest precision.

To better characterize pre-analytic factors that may be affecting TL assay precision, we tested how commonly utilized methods for assessing DNA quality, including purity, and integrity predicted measurement precision as assessed in our study. Recognizing that a myriad of different aspects may contribute to variation in enzymatic processes, including primer sequence (e.g. single copy gene), primer binding, Taq polymerase efficiency, etc., it was important to leverage the existing study design and evaluate current metrics as a first step in determining what factors contribute to variability in precision. Nevertheless, we find that all three quality control metrics, DNA integrity (DIN), purity (OD260/230 and OD260/280 ratios) predicted measurement precision to a non-negligible extent. However, while low scores on these metrics predict low measurement precision in most cases, high scores yielded high as well as low measurement precision. Variation in OD260/230 ratio within batches was consistently positively associated with variation in measurement precision, increasing confidence in the value of this metric of sample quality with relevance to qPCR-based method precision. In contrast, within batch variation in the OD260/280 ratio was not associated with variation in measurement precision, suggesting that the association between the OD260/280 ratio and measurement precision on the batch level can be attributed to another batch level factor that independently affects both the OD260/280 ratio and measurement precision. (18, 22).

Despite the rigorous scientific design, limitations remain. We only compared DNA extractions across singleplex qPCR-based and Southern blot and did not evaluate multiplex qPCR-based methods, nor other methods such as STELA, TeSLA, Luminex, Flow-FISH, etc. While our current conclusions are only applicable to singleplex qPCR-based TL assays, the TRN is currently conducting additional studies incorporating aTL, MMqPCR, TL derived from DNAm data, nanopore sequencing, and additional laboratories performing Southern blot (https://trn.tulane.edu/network-activities-opportunities/method-comparison-studies/). One laboratory received samples that during shipping thawed, an unfortunately not uncommon experience for large scale population-based studies. Although the DIN from these samples appeared to be lower, we were under-powered to statistically test the effect on the ICCs directly. Our findings are also limited to effects in whole blood. Whether similar differences in assay precision across DNA extraction protocols will be found with PBMCs or other non-invasive or minimally invasive samples such as DBS, saliva or buccal swabs remains to be tested. We selected the most commonly utilized extraction methods but were limited in the number of different extraction protocols as well as the number of labs performing each extraction. Previous studies have reported that the effect of DNA extraction varied even within the DNA extraction kits using the same underlying principle (23). There are several approaches to determining assay precision and this includes both internal (within laboratories) and external (across methods/laboratory) comparisons. We share previous concerns that given the established range of factors able to affect qPCR TL assay precision, efforts to regularly confirm internal assay precision, potential integration of external validation practices, and, particularly for new laboratories establishing the methodology, exhaustive and repeated efforts to ensure TL assay precision is paramount to ensuring scientific rigor for TL studies of any size (24).

It is notable that while TRF measurement of repeat extractions has a higher ICC, TL values derived from TRF may intrinsically vary from TL measured by qPCR due to assay differences in the potential for measurement to include variable amounts of sub-telomeric regions and interstitial telomeric sequences. There is a clear need to better understand how TL measurement, using any assay, is impacted by polymorphic variation in subtelomeric regions and interstitial telomeric repeats. The use of direct sequencing via nanopore technology offers the unique possibility of generating a true “gold” standard TL measurement (25). Further, the completion of the sequencing of the human genome, telomere to telomere, is an opportunity to define the unique DNA sequences bridging the subtelomeric region and the telomere for each chromosome arm and leverage existing technologies such as STELA to determine the specific length of each unique chromosome (26). The development and/or validation of a gold standard measure of TL able to generate an absolute length, in base pairs, of TL without inclusion of variable subtelomeric sequences and, ideally, also able to measure TL across the entire spectrum of lengths in humans, is a key goal of the TRN.

## Conclusions

Understanding the importance of pre-analytical factors on the precision of TL assays has significance for the multitude of studies interrogating TL as an indicator of exposure or predictor of disease risk. Our study is the most scientifically rigorous to date with blinded analyses of duplicate extractions and duplicate TL measurement across three distinct DNA extraction types and multiple laboratories. Although we are unable to state which DNA extraction method is “best” for TL measurement, we are able to confirm that even within laboratories which regularly perform TL measurement there is significant inter- and intra-assay variability related to DNA extraction. Our data provides unexpected evidence that the ICCs of measurement on the same extracted DNA sample is not a reliable indicator of measurement precision. When calculating power for epidemiologic studies we have previously reported sample size estimates based on ICCs(1). However, we are now better positioned to recommend that the ICC be calculated based on TL measurement of independent DNA extractions of the same sample and time-point for sample size calculation. For longitudinal studies of TL, the impact of lower ICCs on power is substantial. We therefore encourage laboratories to calculate, and report, ICCs based on (at least) two independent DNA extractions for all studies on a minimum of 30 samples, or at least 10% of the total samples as the strongest indicator of assay precision and demonstrate the influence of independent measurement on ICCs for additional guidance (S6 Figs). Individuals reviewing results and/or funding applications should ensure opportunities for complete methodologic reporting as another way to ensure scientific rigor. Our results indicate important limitations to TL measurement utilizing qPCR-based approaches. Despite these limitations, the existing substantial literature supported by meta-analysis and the recent results from the UK biobank study provide powerful evidence of the utility of q-PCR based methods for adequately powered population-based studies that rigorously evaluate, and report, assay precision (27, 28). Advancing the mechanistic understandings of TL for human health and aging will also require clarification of the specific aspect of telomere length, e.g. distribution of TL, amount of short telomeres, presence of critically short telomeres, or chromosome specific lengths, as the most predictive of exposures or human disease. Moving forward, in addition to ensuring adequate statistical power, future studies will continue to need to balance practicality, precision, and the specific characteristic of TL predicted to relate to the proposed biological pathway.

## Acknowledgements

Thank you to the members of our external advisory committee (Daniel Belsky, Jerry Shay, Daniel Nettle, Dan Eisenberg, Sonja Entringer, and Eileen Crimmins) as well as our NIH partners (Max Guo, Michelle Heacock, and Lisbeth Nielsen for their continuous support of the Telomere Research Network activities and U01 laboratories.

## Supporting information

**S1 Fig.**
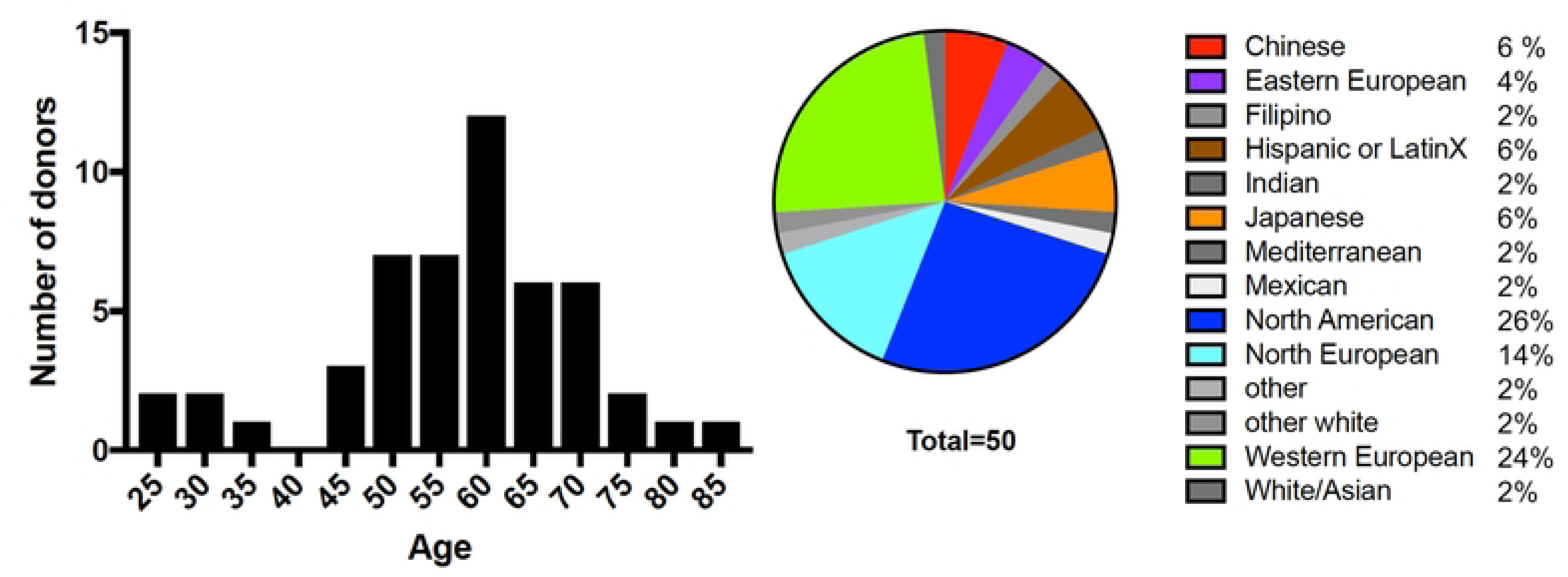
Age and ethnicity distribution of the 50 donors.

**S2 Fig.**
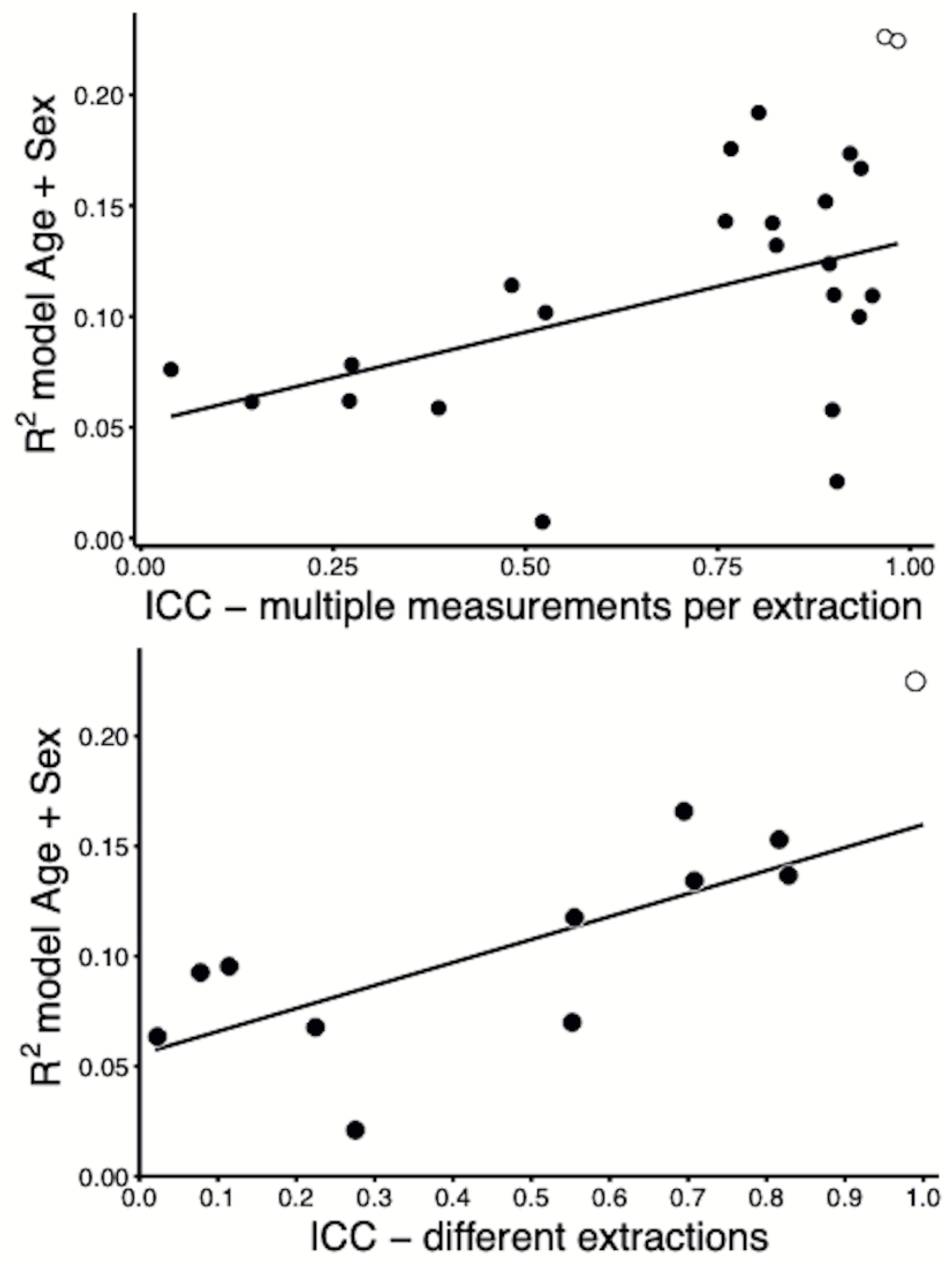
Variance explained (R^2^) in telomere length by age and sex (combined in one model) in relation to qPCR measurement precision. A. Intraclass Correlation Coefficient, calculated over measurements of a single extraction of each sample (R^2^ = 0.23, calculated excluding the Southern blot results). B. ICC calculated over measurements of two extractions of each sample (R^2^ = 0.48, calculated excluding the Southern blot results). qPCR: solid dots; Southern blot: white dots.

**S3 Fig.**
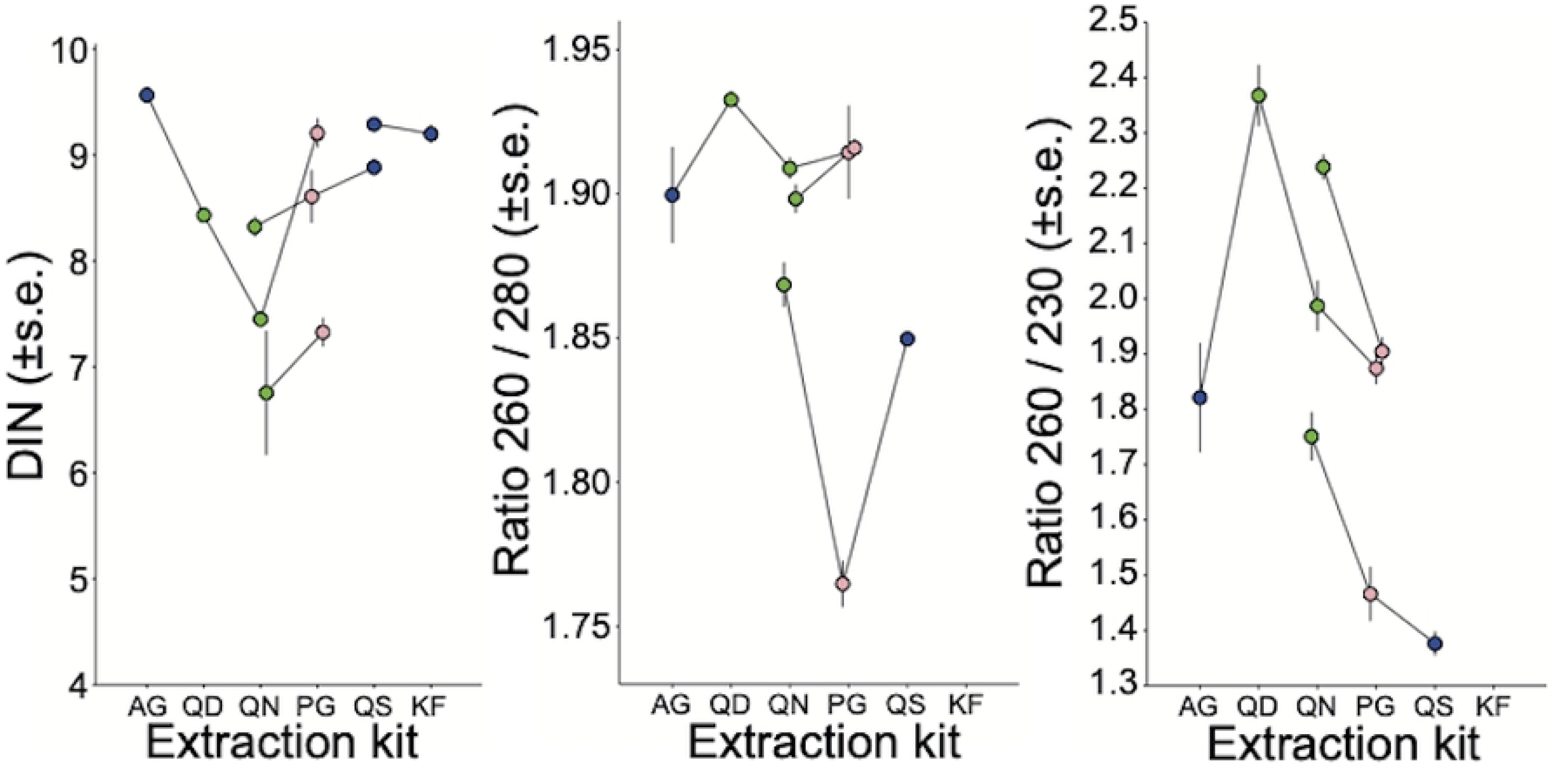
Correlation of DIN, 260/280, and 260/230 across repeated extractions and labs.

**S4 Fig.**
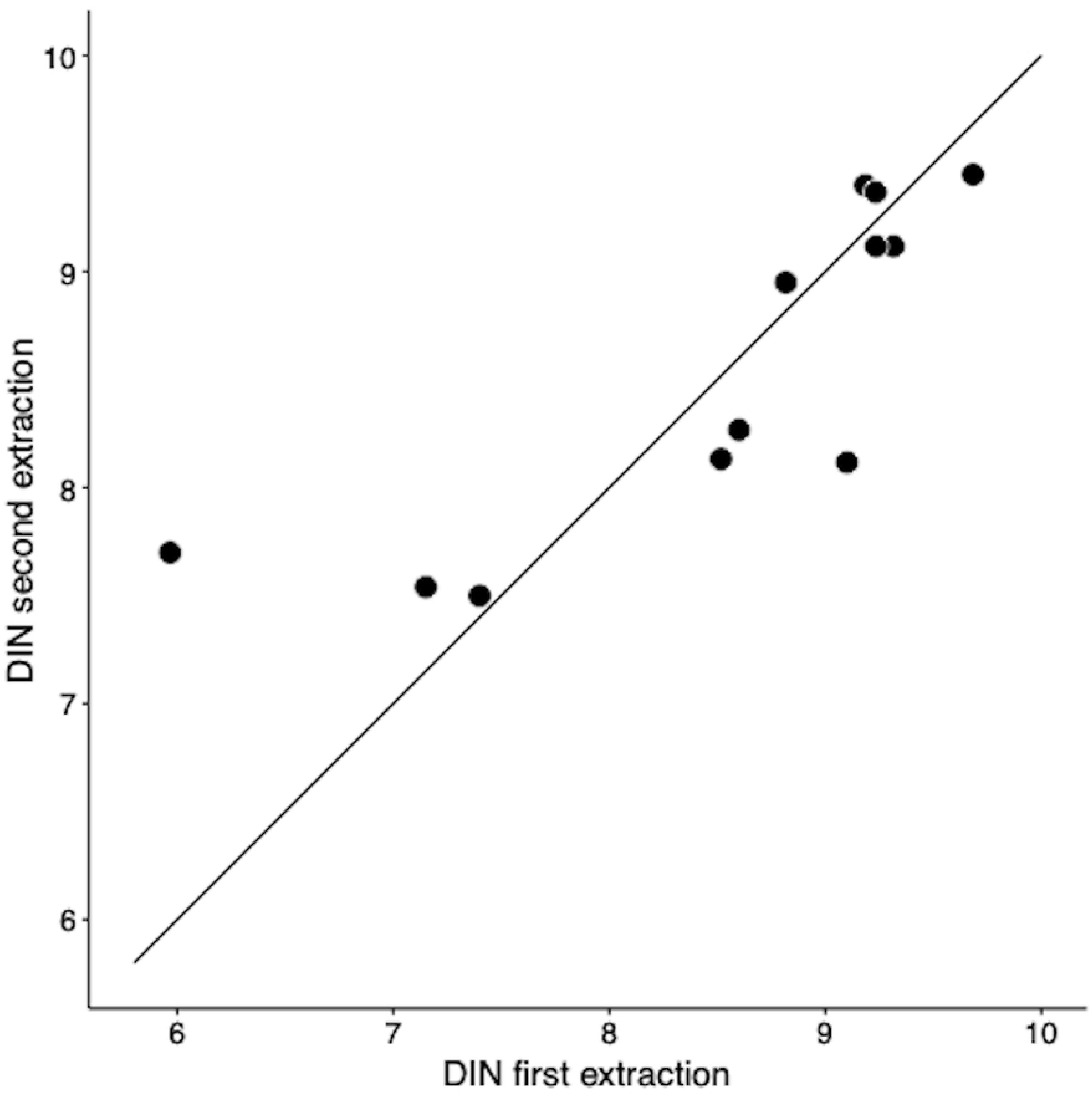
Repeatability of DIN.

**S5 Fig.**
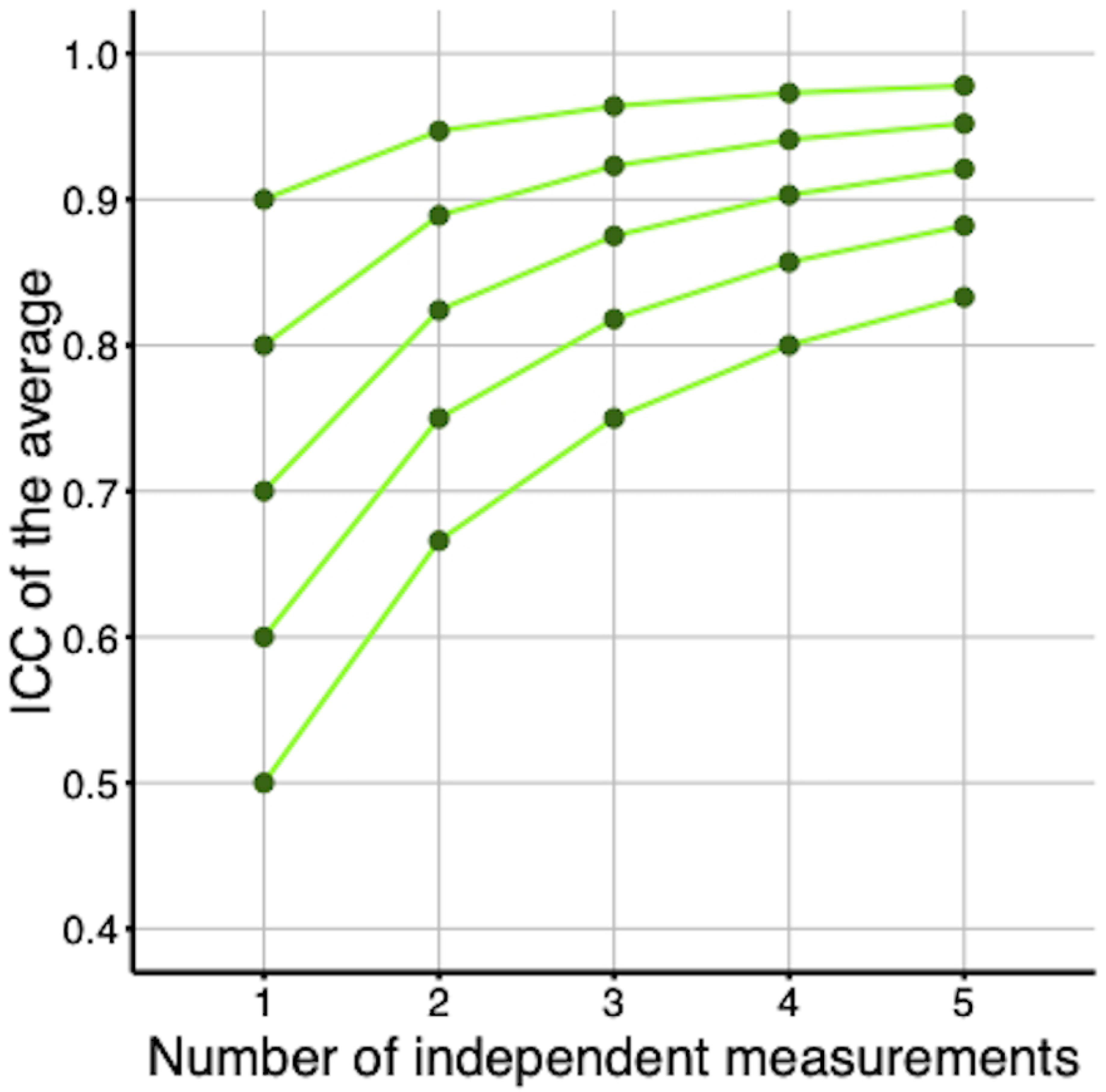
Quality control metrics in relation to extraction kit and extraction principle. A. DIN, B. OD260/230, C. OD260/2880.

**S6 Fig.** ICC based on independent measurements.

**S1 Protocol. Lab 1 telomere length measurement protocol.**

**S2 Protocol. Lab 2 telomere length measurement protocol.**

**S3 Protocol. Lab 3 telomere length measurement protocol.**

**S4 Protocol. Lab 4 telomere length measurement protocol.**

**S5 Protocol. Lab 5 telomere length measurement protocol.**

